# Predictive whisker kinematics reveal context-dependent sensorimotor strategies

**DOI:** 10.1101/833103

**Authors:** Avner Wallach, David Deutsch, Tess Oram, Ehud Ahissar

## Abstract

Animals actively move their sensory organs in order to acquire sensory information. Some rodents, such as mice and rats, employ cyclic scanning motions of their facial whiskers to explore their proximal surrounding, a behavior known as whisking. Here we investigated the contingency of whisking kinematics on the animal’s behavioral context that arises from both internal processes (attention and expectations) and external constraints (available sensory and motor degrees of freedom). We recorded rat whisking at high temporal resolution in two experimental contexts - freely moving or head-fixed – and two spatial sensory configurations – a single row or three caudal whiskers on each side of the snout. We found that rapid sensorimotor twitches, called pumps, occurring during free-air whisking carry information about the rat’s upcoming exploratory direction, as demonstrated by the ability of these pumps to predict consequent head and body locomotion. Specifically, pump behavior during both voluntary motionlessness and imposed head-fixation exposed a backward redistribution of sensorimotor exploratory resources. Further, head-fixed rats employed a wide range of whisking profiles to compensate for the loss of head- and body-motor degrees of freedom. Finally, changing the number of intact vibrissae available to a rat resulted in an alteration of whisking strategy consistent with the rat actively reallocating its remaining resources. In sum, this work shows that rats adapt their active exploratory behavior in a “homeostatic” attempt to preserve sensorimotor coverage under changing environmental conditions and changing sensory capacities, including those imposed by various laboratory conditions.

## Introduction

Perception is a process in which the sensory organ is actively employed in order to acquire sensory data from the external environment (1–6). In his classic study, Alfred L. Yarbus (1) demonstrated active sensing in human visual perception; Yarbus showed that different behavioral contexts, determined by giving subjects perceptual instructions, entail different spatial sampling strategies. A similar approach was employed to study sensorimotor exploration in other visual animals (7, 8), yet little is known about the effects of context on spatial sampling in other modalities.

Many mammals use the long hairs (vibrissae or whiskers) on either side of their snout to navigate the environment, and to collect information about their proximal surroundings (9). In some rodents, movements of the whisker array are used to actively acquire tactile information about both the position and nature of nearby objects (10). These movements are, in turn, affected by the acquired sensory information, as well as by other ‘top-down’ modulatory processes (11–13). In other words, perception is not solely active, but is also reactive, giving rise to closed-loop dynamics of the perceiving organism and its environment (14–16).

Several studies have described basic components of vibrissal active behavior, both those observed in synchronous exploratory whisking in air (2, 17–21) and those related to interactions with external objects (22–25). Only a handful of studies, however, have analyzed how vibrissal behavior is affected by behavioral context. Arkley and colleagues (26), for instance, showed dramatic effects of training and environmental familiarity on the whisking strategy employed by rats. Furthermore, they showed that whisking strategy reflects the animals’ expectation of future object encounters. In another study by the same group, Grant and colleagues (27) tracked the developmental emergence of previously described behaviors in the pups’ first post-natal weeks. Finally, rats change their whisking strategy in response to external perturbations, keeping some behavioral variables controlled, while others are modulated in order to maintain perceptual performance (28). The effects of behavioral context on free exploratory whisking, however, remain poorly described.

In laboratory experiments, highly dominant contextual factors emerge from the experimental methodology. Experimental biologists are often forced to impose methodological constraints on their study subjects to ensure precise and stable observations that are amenable to analysis. One of the most common practices in neurophysiological and neuroimaging studies is *motion restraint*, examples of which are (i) head-fixing, where head movements are eliminated by the physical anchoring of the head (29); and (ii) body restraint, where head movements are permitted while the body is restrained (21). Such procedures entail drastic reduction of the motor degrees of freedom available to the animal, as well as the introduction of psychological stress (29). An additional practice which is prevalent in the study of vibrissal perception is the reduction of the number of vibrissae available to the rodent, either by trimming or by plucking them from a full pad of ~33 macrovibrissae that are arranged in 5 rows to (i) a single row (30), (ii) a few whiskers (23), or (iii) none at all (9); in many cases this is done to facilitate precise measurement of whisker position and shape using overhead videography. This procedure directly and selectively reduces the rodents’ sensory degrees of freedom, and was shown to entail compensatory behavioral adjustments in the context of object interrogation (31). Despite the ubiquity of such manipulations and their possible implications on the motosensory system, no attempt has been made so far to quantify the adaptations they might entail during exploratory behavior.

Here we compare different aspects of vibrissal behavior measured in three contexts (**Fig 1A**): (i) head-fixed rats with a trimmed whisker pad (in this case, only three caudal macrovibrrisae - C1, C2 and D1-were untrimmed on either side; reuse of data published in (23)), (ii) freely moving rats with the same three-whisker configuration as in (i), and (iii) freely moving rats with an entire single row (row C) of macrovibrissae in either side (termed here ‘free single-row’, for the sake of brevity; behavioral apparatus is illustrated in **Fig S1**). We focused only on segments in which the animals performed exploratory rhythmic movements in free-air, without encountering any object with the whiskers. It should be noted that whisker contacts with the floor couldn’t be ruled-out in the freely moving rats due to video limitations (resolution and focus). However, based on a previous study (13) we can estimate the probability of such floor contacts to be only 2.5% per cycle for any whisker and therefore are not expected to significantly alter our findings. Moreover, headfixed rats were positioned such that no floor contacts were possible. We begin by describing the different parameters of whisking behavior used in our analyses, and then focus our analysis on a kinematic feature called the *free-air pump* (20, 32). We then show that headfixing exerts a dramatic shift in the whisking profile, suggesting an adaptive redistribution of sensorimotor exploratory resources. Finally, we demonstrate that whisker trimming likewise entails an alteration in whisking strategy.

**Fig 1:**
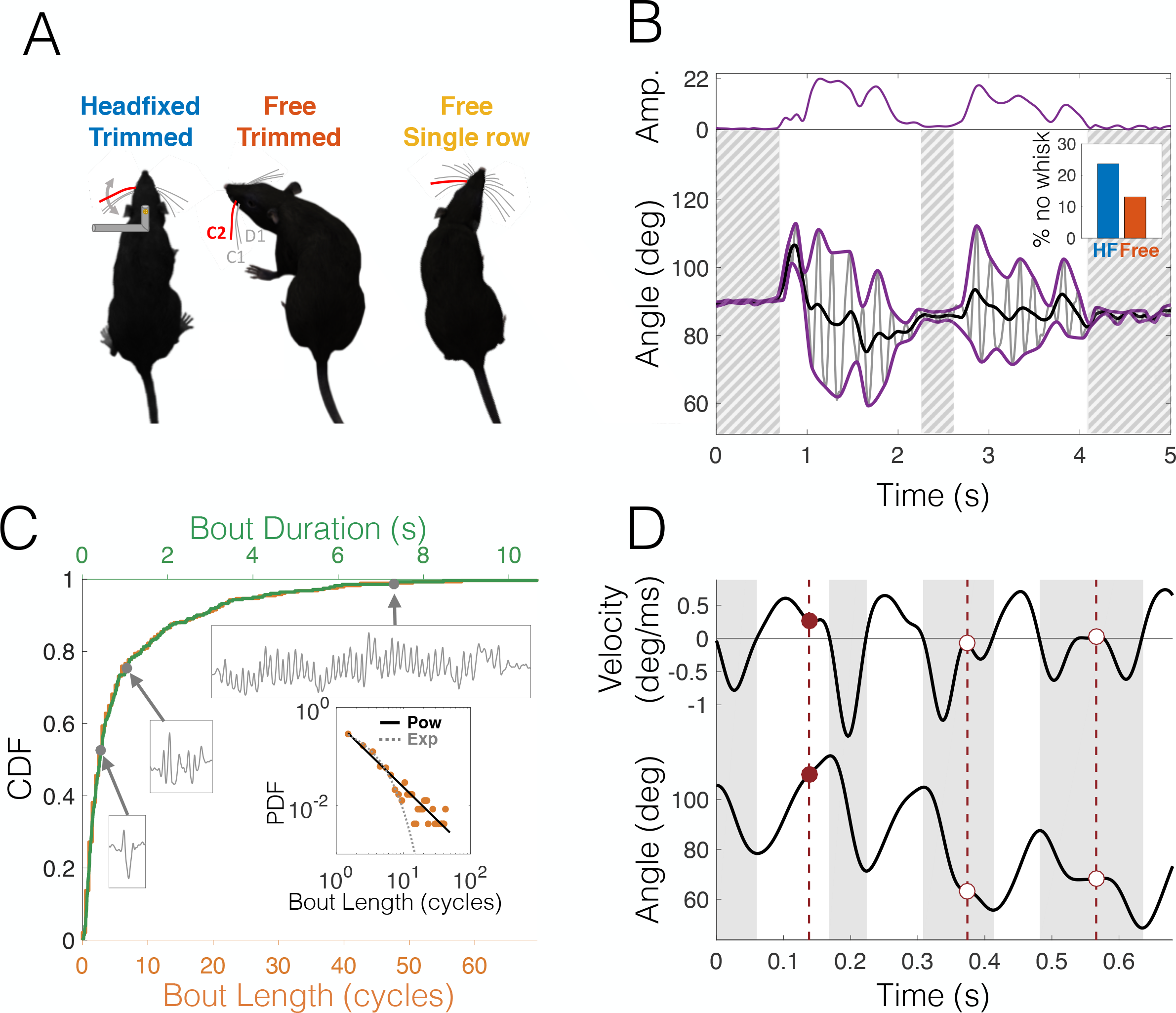
Methodology, whisking bouts and free-air pumps. **(A)** Three datasets were used in this study: headfixed rats with only whiskers D1, C1 and C2 (left); freely-moving rats with only whiskers D1, C1 and C2 (free trimmed, center); and freely-moving rats with 7 whiskers of row C (C1-C7) (free single row, right); only motion of C2 (in red) was used in all analyses; only left side whisker was used in all analyses except in Figs 3 and 4. **(B)** Rhythmic decomposition of whisking. Protraction angle (grey line) was analyzed as fast oscillations modulated by multiplicative amplitude (top, purple) and additive offset (black). Hatched: non-whisking epochs (amplitude<2 deg). Inset: headfixed animals spend less time whisking. **(C)** Cumulative distribution function of bout duration in cycles (bottom abscissa, orange) and in seconds (top abscissa, green) in headfixed rats. Examples of bouts 0.44 s, 1.04 s and 7.3 s long are shown. Inset: Probability Density Histogram of bout length is heavy-tailed; Solid black line: power-law fit, dotted line: best exponential fit, shown for comparison. **(D)** Example of a whisking trajectory. Bottom: protraction angle; top: angular velocity. White background: protraction phases; grey background: retraction phases. Red markers and vertical dashed lines: free-air pumps, phases in which velocity profile is double-peaked; filled circle: protraction pump, empty circles: retraction pumps.

## Results

### Characterizing whisking behavior

All analyses were performed on the protraction angle of whisker C2 (protraction angles of different whiskers are highly correlated during exploratory free-air whisking, (20)). In the analysis of head-motion (Figs 3 and 4) we used C2 of both sides of the face; in all other analyses, only the left side was used. To quantify the governing variables of exploratory whisking, we employed ‘rhythmic decomposition’ on each of the tracked segments (21); this algorithm models whisking dynamics as rapid oscillations (whisking cycles), modulated by a multiplicative *amplitude* process and an additive *offset* process (see **Fig 1B** and Methods). The whisking process consists of *bouts* (amplitude > 2 deg, non-shaded in **Fig 1B**) separated by stretches of non-whisking (amplitude ≤2 deg, hatched in **Fig 1B**); the 2 deg threshold was set based on the bimodal distribution of whisking amplitudes (see **Fig S2**). Overall, head-fixed rats whisked less frequently than freely moving, with head-fixed rats whisking 76.4% of the tracked time, while freely moving rats whisked 89.8% of the tracked time (inset, **Fig 1B**). This reflects the well-known reluctance of head-restrained rats to whisk; often, some sensory stimulation (e.g., olfactory) is required to encourage head-fixed animals to whisk. The duration of whisking bouts varied greatly (**Fig 1C**); in the head-fixed dataset, the median bout duration was 0.412 s (2.5 whisking cycles) while the mean was 0.973s (6.2 cycles) (N=283). This large difference between median and mean reflects the ‘heavy tail’ of the bout distribution. Indeed, the bout probability density is well fitted with a power-law function (inset of **Fig 1C**). The freely-moving datasets did not contain enough complete bouts (where both the beginning and the end of the bout are recorded) to allow for a statistical analysis. All analyses described below were performed on bout epochs (i.e., non-whisking epochs were excluded).

Each whisking cycle is comprised of two stages: protraction, in which the whiskers move rostrally, and retraction, in which they move caudally (unshaded and shaded in **Fig 1D**, respectively); henceforth we will refer to protractions and retractions as the two *phases* of the whisking cycle. It was noted in several previous studies that the velocity profile of individual phases is occasionally multi-peaked, a feature termed ‘pump’ (red markers in **Fig 1D**(20, 32)). The occurrence of these pumps in the context of object contact (Touch Induced Pumps or *TIPs* (23)) was shown to be related to object-oriented spatial attention (13). Next we compare various properties of ‘*free-air pumps*’ and TIPs.

### Free-air pumps are temporally clustered and entail phase prolongation and offset shifting

TIPs cluster in time (23). We tested the tendency of free-air pumps to cluster by measuring the cross-correlation of pump instances between cycles. The probability of free-air pump occurrence was significantly correlated across multiple cycles (**Fig 2A_1,2_**). Importantly, these correlations were specific to pumps of the same whisking phase: a cycle in which a pump occurred in the protraction phase was likely to be preceded and followed by other cycles with protraction-pumps (p<0.05 up to lags of 7 cycles, random permutations, **Fig 2A_1_**), and the same was observed for retraction pumps (p<0.05 for up to and beyond 10 cycles, random permutations, **Fig 2A_2_**). Sequences of consecutive protraction/retraction pumps were significantly more frequent than those obtained by random permutations (p<0.0002 and p=0.0006, circles, **Fig 2B_1,2_**) and the same was true for sequences of no-pumps (p=0.025 and p=0.0022, squares, **Fig 2B_1,2_**). Retraction pumps were less likely to be immediately followed by protraction pumps (p<0.005, random permutations, grey arrowheads in **Fig 2A_1,2_**), but otherwise pumps of opposing phases were uncorrelated.

**Fig 2:**
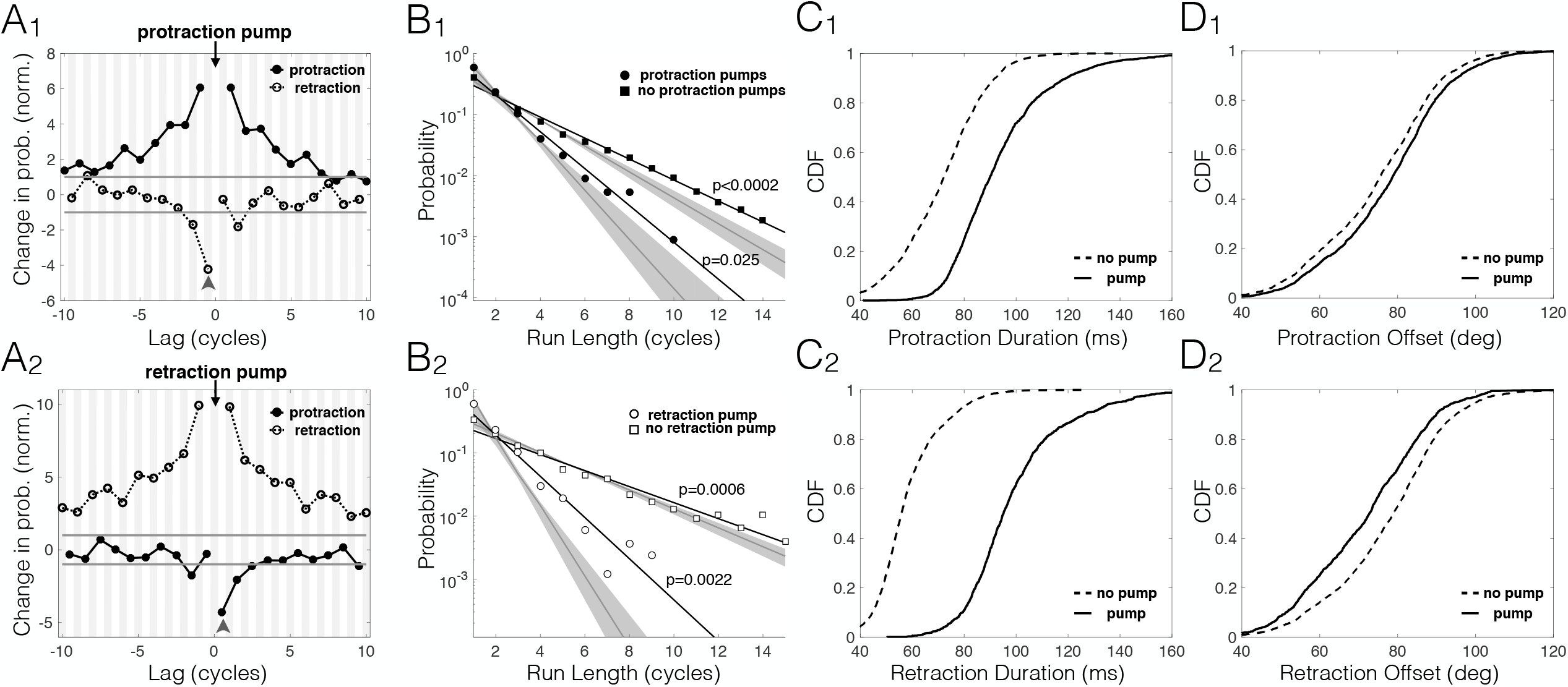
Phases with free-air pumps are clustered in time. **(A_1,2_)** Temporal clustering. Change in pump probability (normalized to the 0.05 significance level, i.e. −1 is the 5^th^ percentile and -1 is the 95^th^ percentile, both marked with a grey horizontal line) in protractions (solid, filled circles, white background) and retractions (dashed, empty circles, grey background), given that a pump (black arrow) occurred in protraction (A_1_, top) or retraction (A_2_, bottom) of a cycle, as function of time lag from that cycle. Same-phase pumps are strongly correlated across multiple cycles. Different-phase pumps are overall uncorrelated, though a retraction pump is unlikely to be immediately followed by a protraction pump (grey arrowheads). **(B_1,2_)** Probability distributions of series of consecutive pumps (circles) and no-pumps (squares) of different run-length; shown separately for protractions (filled markers, B_1_, top) and retractions (empty markers, B_2_, bottom). Long sequences of pumps/no-pumps are significantly more common than chance level due to the temporal correlations shown in A. Grey shading: 0.95 confidence margins for controls generated by random permutations. **(C_1,2_)** Phase prolongation. Cumulative density function of phase duration for protractions (C_1_, top) and retractions (C_2_, bottom) with pumps (solid) and without pumps (dashed). Phases with pumps are longer (means: protractions 93 ms/71 ms, retractions 99 ms/58 ms with/without pumps, respectively). **(D_1,2_)** Offset shifting. Cumulative density function of mean whisking offset of protractions (D_1_, top) and retractions (D_2_, bottom) with pumps (solid) and without pumps (dashed). The offset during phases with pumps are shifted towards the direction of motion, i.e. forward in protractions and backwards in retractions (means: protractions 77.7 deg/74.6 deg, retractions 77.3 deg/71.5 deg with/without pumps, respectively).

**Fig 3:**
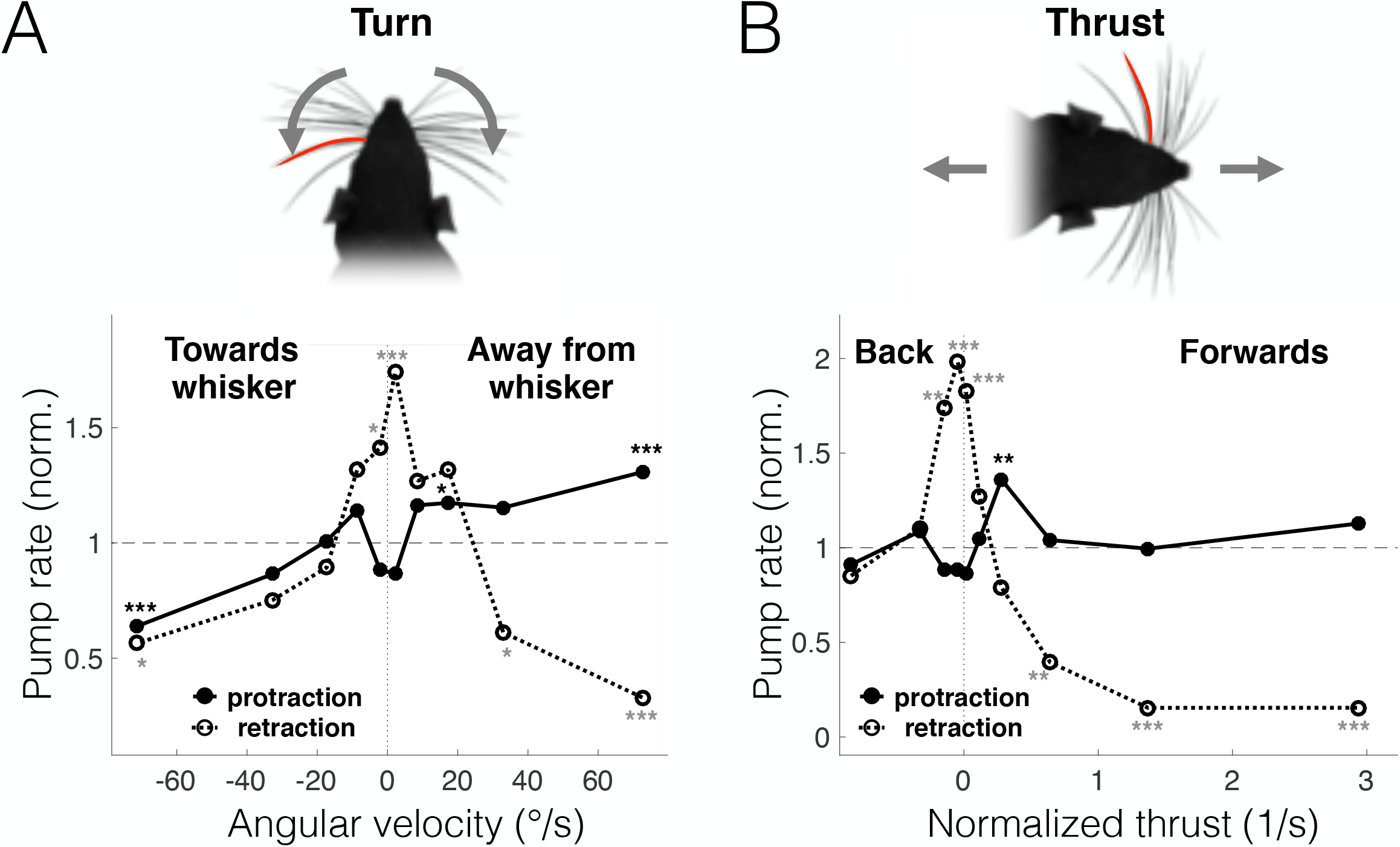
Free-air pumps are related to head motion. Pump rate (normalized to mean pump rate across all cycles) conditioned of head-motion variables in freely moving rats. Motion of whisker C2 from both sides of the snout was used in these analyses. Protraction: solid line with filled circles; retraction: dashed line with empty circles. **(A)** Turns. Protraction pumps are more frequent in contralateral turns, i.e. when the pumping-whisker side moves forward. Retraction pumps are more frequent when the rat does not turn its head. **(B)** Thrust. Protraction pumps are more frequent in low-velocity motion forward, while protraction pumps are more frequent when the rat is close to stationary. All linear velocities are normalized to head size. *=p<0.05; **=p<0.01; ***=p<0.005; no symbol=not significant.

**Fig 4:**
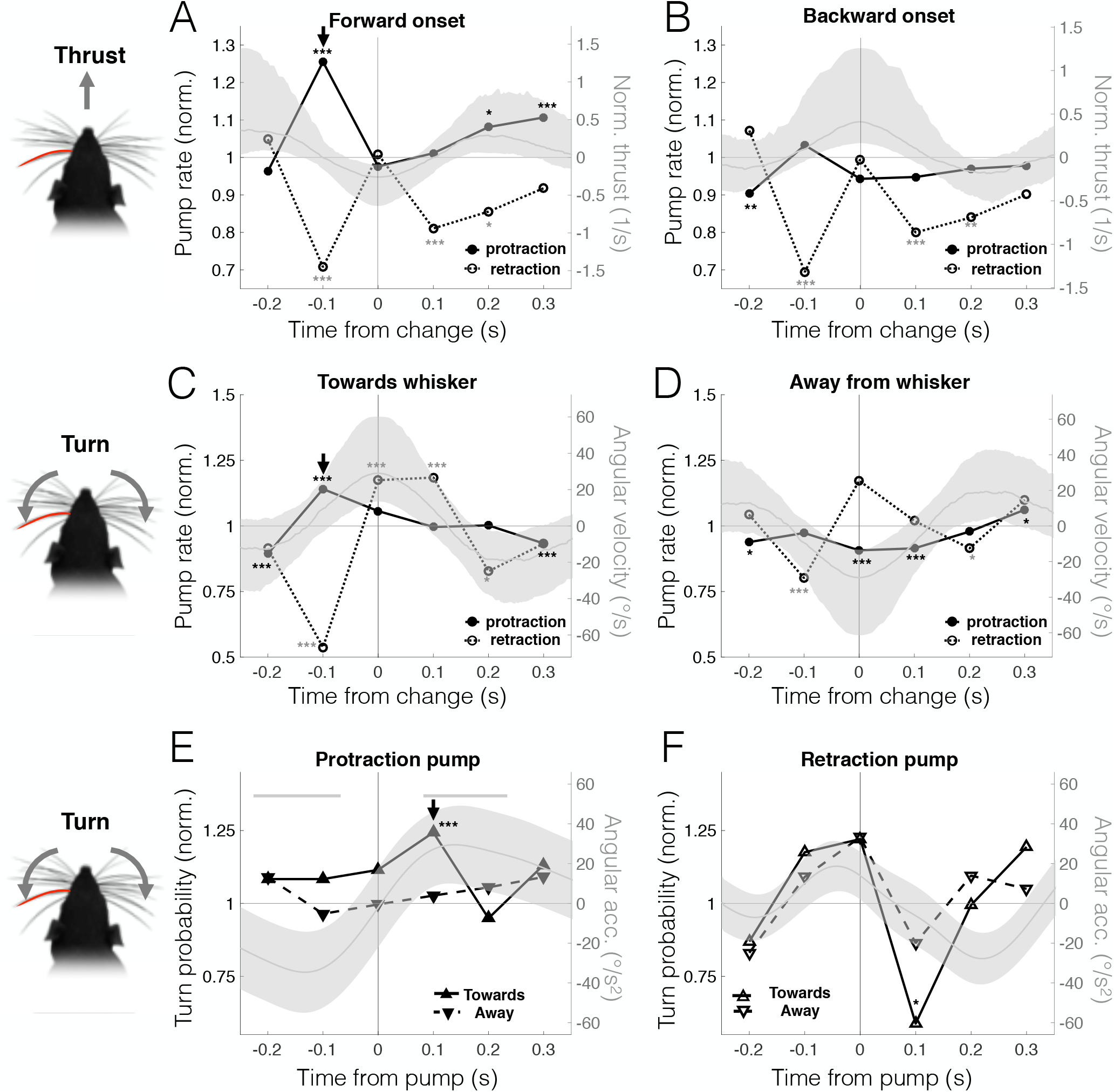
Free-air pumps predict head motion. Dynamics of pump rate (normalized to mean rate) aligned to onset of head motion (i.e., zero-crossing of acceleration). All probabilities were calculated in 100 ms wide bins. **(A-B)** Protraction pump rate (left ordinate) is significantly elevated 100 ms prior to onset of forward motion (p<2·10^−4^, random permutations). No such increase is present in onset of backward motion (p=0.22, random permutations). Right ordinate: thrust (normalized to head size); median-dark grey curves, inter-quartile range-shaded grey area. **(C-D)** Protraction pump rate (left ordinate) is significantly elevated 100 ms prior to head turn toward pumping whisker side (p<2·10^−4^, random permutations). No similar increase is present in onset of turn away from pumping whisker side (p=0.26, random permutations). Right ordinate: angular velocity; median-dark grey curves, inter-quartile range-shaded grey area. **(E-F)** Ipsilateral turns probability (left ordinate) is elevated 100 ms after the occurrence of a protraction pump (p=, random permutations), but depressed 100 ms after a retraction pumps (p=0.02, random permutations). Right ordinate: angular acceleration; median-dark grey curves, inter-quartile range-shaded grey area. Thick grey line: statistically significant times were average acceleration was non-zero (p<0.05, random permutations). *=p<0.05; **=p<0.01; ***=p<0.005; no symbol=not significant.

Did cycles containing free-air pumps differ from those lacking them? In the context of object encounters, protractions containing TIPs are longer and shifted forward when compared with protractions with no pumps (23). This was also true for free-air pumps: first, phases with such pumps were much longer than those lacking pumps (31% longer in protraction and 71% longer is retraction, **Fig 2C_1,2_**, as previously shown in (20)). Second, the whisking offset (i.e., the midpoint angle of the motion) was shifted in the direction of the whisking phase that included the pumps, i.e. more protracted in protractions that included pumps and more retracted in retractions that included pumps (+4.2% and −7.6% change in protraction/retraction, respectively, **Fig 2D_1,2_**). We note, however, that the statistical significance of this last finding is difficult to assess due to temporal correlations in the offset and pump-rate signals, which render neighboring samples statistically dependent. Taken together, we conclude that free-air pumps during exploratory whisking exhibit the key temporal (clustering and duration) and spatial (offset) characteristics of the TIPs, which were previously shown to be related to object-oriented spatial attention.

### Direction specific correlations between free-air pumps and head motion

We decomposed head motion into three components: *turn* (rotation around the midpoint between the eyes), *thrust* (longitudinal translation or forward/backward motion) and *slip* (transverse translation or side motion). While turning was uncorrelated with thrust (Pearson coefficient R=−0.035), it was highly correlated with slip (Pearson coefficient R=0.863, see **Fig S3**); these correlations reflect the fact that the axis of head rotation (the neck) is caudal to the eyes (which were the feature tracked in our analysis).

Therefore, we limit our analysis of head motion to the turn and thrust variables. Protraction pumps were significantly more frequent during turns contralateral to the pumping whisker, when that whisker’s side moved forward (e.g., pumps of the left whisker were more common during turns to the right, **Fig 3A**) and during forward thrust at moderate speeds (i.e., walking but not running, **Fig 3B**). Correspondingly, protraction pumps were scarce during turns ipsilateral to the pumping whisker, when the whisker side moved backwards. In contrast, retraction pumps occurred mostly when the rat kept its head motionless (close to zero velocity of turn and thrust), and were significantly inhibited during rapid motion in any direction. Therefore, head motion was accompanied with direction-specific modulations in the occurrence of free-air pumps; protraction pumps were mostly directed towards newly explored space, whereas retraction pumps were associated with lingering in one spatial location while keeping both head and body motionless.

### Free-air pumps are predictive of changes in head motion

We next checked whether free-air pumps are linked to changes in the animal’s exploratory behavior. We performed event-triggered analysis to explore the reciprocal relations between head and pump dynamics. First, we identified motion-change events, where the rat changed the direction of head turning or thrust. We then measured the dynamics of free-air pump rate, relative to the onset of the change (the onset of acceleration in the opposite direction). Onset of forward motion was preceded by a substantial increase in protraction pumps, peaking around 100 ms prior to the change (p<2·10^−4^, random permutations, black arrow, **Fig 4A**). No such change occurred in the onset of backward motion (p=0.22, random permutations, **Fig 4B**). Note that in both cases retraction pumps were inhibited before and after the change occurred. Similarly, ipsilateral head turns (*towards* the pumping whisker side) were *preceded* by an increase in protraction pump rate (p<2·10^−4^, random permutations, black arrow, **Fig 4C**), while no such increase is seen prior to contralateral turns (p=0.26, random permutations, **Fig 4D**).

The predictive power of free-air pumps regarding future spatial targets of the rat is confirmed by measuring the dynamics of head turning probability relative to the time a pump occurred. Consistent with the previous analysis, the average protraction pump was followed by an increased probability of turning towards the pumping whisker side (p= 0.0038, random permutations, left ordinate in **Fig 4E**), which is also evident in the dynamics of angular acceleration (p>0.05, random permutations, right ordinate in **Fig 4E**). Retraction pumps, however, were followed by a significant drop in this probability (p=0.02, random permutations, **Fig 4F**). These results are not sensitive to the choice of motion-change detection threshold (see **Fig S4**). We conclude that free-air pumps predict the rats’ spatial motion targets: motion towards newly explored space was preceded by an increase in ipsilateral protraction pumps and a drop in retraction pumps.

### Head-fixed effects on whisking resemble those of voluntary motionlessness

Are voluntary and imposed motionlessness similar in their effects on whisking kinematics? To answer this question, we compared the frequency of protraction and retraction pumps in head-fixed rats with those observed in freely moving rats at different motion speed ranges (**Fig 5A**). When the entire dataset of freely-moving episodes is analyzed (maximal head velocity="∞"), head-fixed rats exhibited 22% fewer protraction pumps than freely moving rats (p=10^−4^, bootstrap, N=4056 and 412 cycles for head-fixed and free trimmed, respectively). In contrast, retraction pumps were dramatically more prevalent in head-fixed rats (208%, p<10^−4^, bootstrap, N=4046 and 549 cycles for head-fixed and free trimmed, respectively). However, as we limit our analysis of pump frequency in freely moving rats to phases in which head velocity did not exceed a certain bound, the frequency of protraction pumps decreases and that of retraction pumps increases to the point of becoming statistically similar (p=0.14, bootstrap, N=75 protractions and 80 retractions), and to the frequencies measured in head-fixed rats (p=0.21 and 0.81 for protractions and retractions respectively, bootstrap, **Fig 5A**). In other words, the overall ratio of protraction-to-retraction pumps drops as velocity decreases, approaching that of head-fixed rats at near motionlessness (**Fig 5B**). A similar trend can be seen in the distribution of protraction/retraction durations (**Fig 5C**). Protractions were significantly longer than retractions for the entire freely moving dataset (means 74 ms and 52 ms, respectively), consistent with previous observations (17, 20, 33–37). However, while both protractions and retractions increased in duration as the maximal speed decreased, the ratio of protraction duration to retraction duration decreased, approaching that of head-fixed rats at near motionlessness (**Fig 5D**). We conclude that freely moving rats, which display a strong emphasis on the protraction phase during motion, approach the protraction-retraction near-parity typical of head-fixed rats as their velocity decreases towards motionlessness.

**Fig 5:**
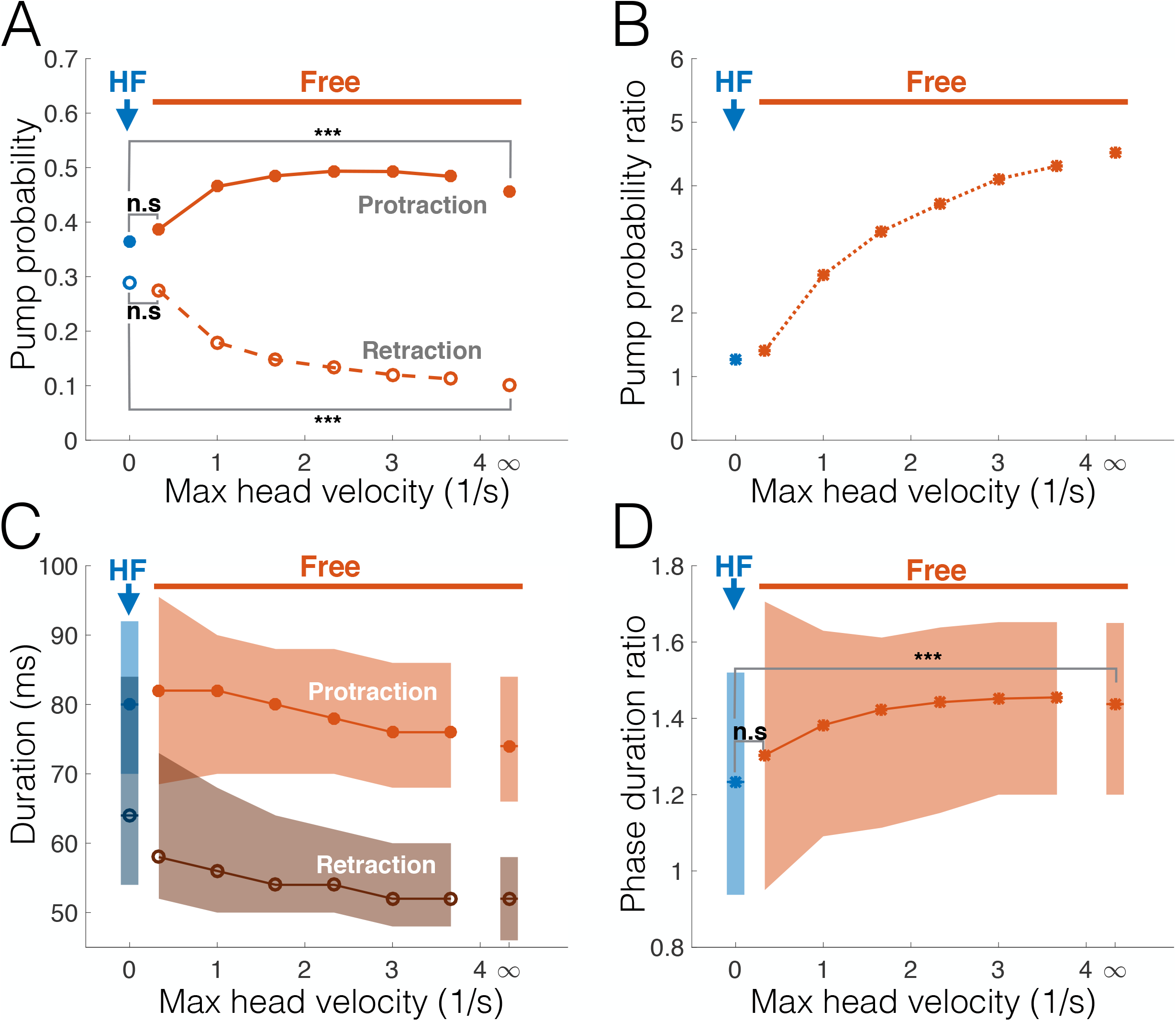
Free-air pumps in headfixed rats resemble those in voluntary motionlessness. Abscissa in all panels: maximal head velocity for samples analyzed from the freely moving dataset (i.e., only cycles in which head-velocity did not exceed this value were taken); ∞=entire dataset. **(A)** Probability of protraction (filled circles) and retraction (empty circles) pumps for headfixed (HF, blue) and freely moving (red) rats. **(B)** Ratio between protraction and retraction pump probabilities. **(C)** Distributions of protraction (median: filled circles; interquartile range: light shading) and retraction (median: empty circles; interquartile range: dark shading) durations for headfixed (blue) and freely moving (red) rats. **(D)** Protraction-retraction duration ratio for headfixed (blue) and freely moving (red) rats (median: asterisks; interquartile range: shading). ***=p<0.005; n.s.=not significant.

### Whisking envelope spatial distribution is dispersed due to headfixing

Our findings so far suggest that limiting the rat’s degrees of motor freedom (by head-fixing) entails a more even distribution of the animal’s sensorimotor resources between protraction and retraction; this was reflected in features such as pump-occurrence and phase duration. Can we see a similar effect in the *whisking envelope* – the combination of amplitude and offset (see **Fig 1B**) that dictates the range of angles covered by whisking at each cycle? It is important to note that while amplitude and offset may be, in principle, uncorrelated, they are necessarily statistically dependent, as the maximal possible amplitude is always dictated by the distance from the current offset to the maximal whisker protraction and retraction angles. The theoretical domain of all possible amplitude-offset combinations, therefore, is bound by an isosceles triangle (**Fig 6**).

**Fig 6:**
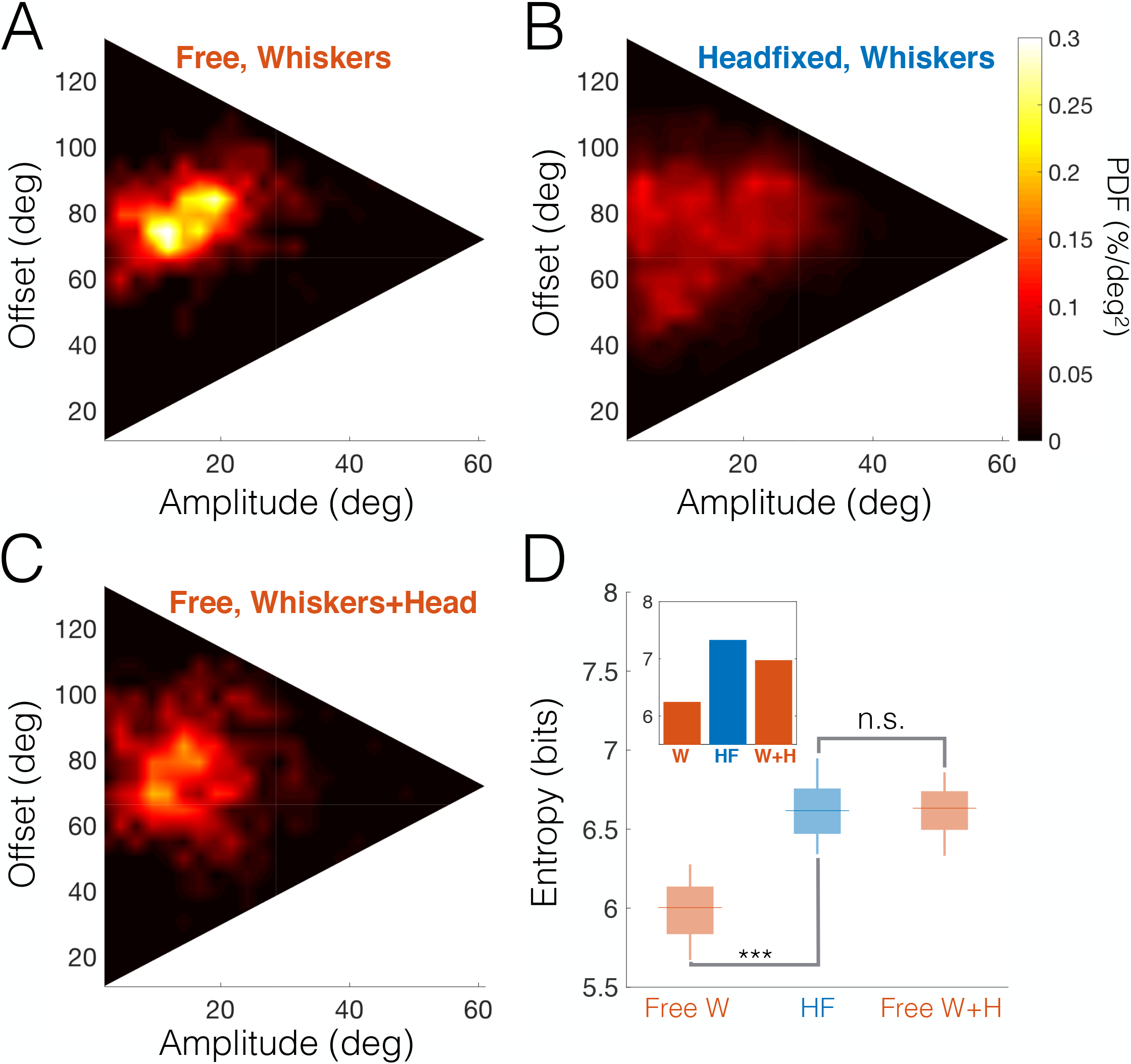
Headfixed whisking envelope distribution compensate for lost degrees of freedom. **(A-B)** Bivariate probability density distributions of whisking amplitude and offset (relative to head) for freely-moving (A) and headfixed rats (B). While freely moving rats exhibit a preferred subspace of amplitude-offset combinations, headfixed rats cover much of the available envelope space (range of offsets is determined by the absolute bounds on whisker C2 angle in all datasets, 11-133 deg; maximal possible amplitude i s offset-dependent, and peaks at the median offset at 61 deg). **(C)** Probability distribution of whisking envelope, taking into account head rotations, for free rats. **(D)** Whisking envelope information entropies for random subsets taken from each dataset (box-and-whisker plots: horizontal line-median; box-IQR). Mean entropy is significantly smaller for freely moving rats (Free W, left) than for headfixed rats (HF, center), indicating that the whisking envelope distribution is much more dispersed during headfixing; however, there is no significant difference when head rotations are included in the analysis (Free W-H, right). ***=p<0.005; n.s.=not significant. Inset: entropies calculated for entire datasets (free whiskers only 6.24 bits, headfixed 7.325 bits, free whiskers-head 6.97 bits).

Utilization of this domain was indeed affected by head-fixing. Comparing rats with the same whisker arrays (head-fixed and freely moving trimmed rats) revealed that while freely moving trimmed rats had a restricted focal region in which they preferentially whisked (i.e., the whisks were narrowly distributed around a preferred amplitude of 11.8 deg and a preferred offset of 74.5 deg, **Fig 6A**), head-fixed rats exhibited a highly dispersed distribution which covered a large portion of the triangular domain (**Fig 6B**). This dispersion may reflect compensation for the lost motor degrees-of-freedom of the head and body. Indeed, whisking envelope distributions became dispersed when changes in head angle were considered in freely moving rats (relative to the mean head angle in each tracked segment, **Fig 6C**). To quantify the envelope dispersion in each scenario, we measured the distributions’ information entropy; to evaluate statistical significance, the bootstrap method was used to generate random subsets of equal size from each dataset (**Fig 6D**, see Methods). The entropy was significantly smaller in freely moving trimmed rats than in head-fixed ones when only whisker angle was used (p<10^−4^, bootstrap), reflecting the increased dispersion of the latter. However, no significant difference was measured when the head’s degree-of-freedom was taken into account (p=0.13, bootstrap). This suggests that the rats compensated for the loss of motor degrees-of-freedom due to head-fixing by employing a wider range of whisking configurations.

### Head-fixing and whisker trimming entail shifts in whisking strategy

We next examined the impact of different experimental constraints on the distributions of individual whisking variables (offset and amplitude, diluted to avoid temporal correlations, see Methods). Comparison of the offset distributions in head-fixed and freely moving trimmed rats (**Fig 7A**) showed that, apart from the increase in variance discussed in the previous section (55% increase, p<10^−4^, bootstrap, N=1294 and 292 for head-fixed and free respectively), head-fixed rats whisked in a slightly retracted angle (4.8% reduction, p=10^−4^, bootstrap). This result further suggests that head-fixed animals explore more retracted positions than free animals, in line with the pump results described above.

**Fig 7:**
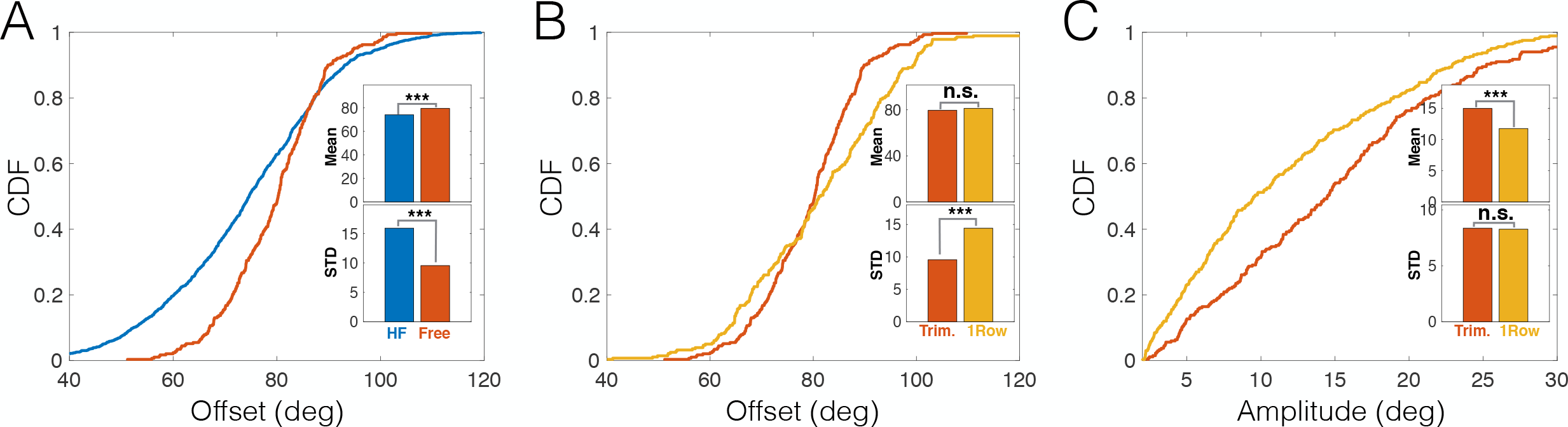
Contextual shifts in whisking strategy. **(A)** Cumulative distribution functions (CDFs) of whisking offset for headfixed (HF, blue) and free (red) trimmed rats. Insets: mean and STD of distributions. **(B)** CDFs of whisking offset of freely moving trimmed (red) and single-row (yellow) rats. Insets: mean and STD of distributions. **(C)** CDFs of whisking amplitude of freely moving trimmed (red) and single-row (yellow) rats. Insets: mean and STD of distributions. ***=p<0.005; n.s.=not significant.

Finally, we analyzed the dependency of the whisking’s spatial pattern on the number of whiskers available to the rat. As described above (see **Fig 1A**), freely moving rats had either a configuration of three caudal whiskers or a full row of seven whiskers (i.e., including both caudal and rostral whiskers). Comparison of the whisking amplitude and offset distributions of these two datasets of freely moving rats revealed that the preferred whisking strategy was noticeably different. The trimming-induced effect consisted of two adjustments: First, a significant 33.8% decrease in the variance of offsets (p<10^−4^, bootstrap, N=292 and 280 for single-row and trimmed, respectively; **Fig 7B**) while maintaining the mean offset almost unchanged (2% decrease, p=0.11, bootstrap); and second, a significant 27.4% increase in mean amplitude (p=10^−4^, bootstrap, N=367 and 268 for single-row and trimmed, respectively; **Fig 7C**) while not significantly changing its variance (p=0.91, bootstrap). Therefore, the trimmed rats tended to employ large amplitude whisks around a relatively constant offset angle, while those having a full row of whiskers used smaller amplitudes while shifting the offset over time.

## Discussion

In this paper, we have shown that whisking kinematics predict consequent head and body locomotion, and, consistently, that these kinematics depend on the behavioral context. The spatial and dynamical characteristics of rat’s exploratory whisking were affected by the rats’ ability to move and the number of whiskers they had available. It is commonly assumed that overt spatial attention is associated with preparing to move the body or the sensors towards a selected location or object (13, 14, 38–41), and therefore we suggest that the alterations in the spatial exploration described here reflect alterations in overt spatial attention. If taken to be true, our findings suggest that both head-fixed rats and free but motionless ones dedicate more attention to whisker retraction than do rats in motion.

This interpretation suggests that the whisking pump, a subtle alteration in the whisking dynamics, is a useful indicator of perceptual attention. It was previously shown that, when induced by encountering an object, such pumps are robustly associated with object-oriented attention (13). While free-air pumps were shown to have different temporal dynamics than those of TIPs (13), suggesting distinct underlying sensory-motor pathways, we demonstrated here that the two types of pumps share several key temporal and spatial features: temporal clustering, phase prolongation, spatial offset shifting and bidirectional relationship with head motion. Critically, we show that free-air pumps predict the rats’ future orienting behavior (compare Fig 1C in (13) and Fig 4E here). The differences in the temporal scales of sensory-motor kinematics (13) can probably be accounted for by differences in the underlying sensory-motor pathways. While TIPs seem to be implemented via brainstem loops (13, 23, 42), free-air pumps may involve higher-order sensory motor loops.

The predictive relations between free-air pumps and the rat’s locomotion may reflect the rat’s expectation of future encounters. Thus, when the rat moves forward at moderate velocity while exploring the environment, it might expect novel encounters to occur mostly during protraction (26), and therefore increases the pump rate in that direction in order to improve sensory acquisition (Fig 3), either by extending protraction duration (Fig 2) or by the pump causing the whisker to briefly ‘revisit’ places of interest. Conversely, when the rat is motionless, either by choice or due to imposed head-fixing, encounters may occur in either direction of whisker motion, and therefore the rat shifts some of its spatial attention towards retraction (Figs 3 and 5). Importantly, this shift in spatial attention was also evident in the overall distribution of the whisking offset in head-fixed rats (Figs 7A).

These findings also offer an interesting way to reconcile an apparent contradiction in previously reported data. Our previous study in anesthetized animals on the sensory representation of whisker motion at the primary afferents and brainstem levels found cells responsive throughout the whisking cycle, with most cells responding to the protraction phase (43). In awake *head-fixed* animals, however, brainstem, thalamic and cortical sensory cells show an overrepresentation of the retraction phase (44, 45). So far, no ethological or physiological explanation was given for this finding. A recent study (46) found correlations between the preferred phase of vibrissal afferents, and the activation of different facial muscle groups controlling whisking motion (47). The facial nerve stimulation method used to evoke whisker motion in anesthetized rats activates mostly one muscle group (the intrinsic whisker pad muscles (47)) involved in protraction, and therefore more cells responding to protraction were sampled. However, we can infer from the results reported here that awake head-fixed animals shift their attention backwards, and may therefore strongly activate the muscle groups involved in controlled retraction (22) and the afferents correlated with that motion. It is also possible that increased attention to retraction involves activation of internal feedback loops within the brain (25, 48) enhancing the activity related to this phase. In other words, the predominance of retraction related cells reported in awake head-fixed animals may reflect the behavioral context imposed by the experimental set-up, rather than the actual distribution of sensitivity in the vibrissal system. The conclusion arising from this possibility is far-reaching: behavioral context may bias physiological findings down to the cellular level, either via alteration of the sensory-motor interactions with the world at the periphery or via internal feedback loops in the central nervous system.

Our freely moving rats displayed a preferred whisking pattern, based on small amplitude whisks around a varying offset. This pattern appears to allow a combination of local active sensation with global (row-wide) passive reception, resembling in part the fovea-periphery division of work in vision. In contrast, trimmed rats applied large amplitude whisks around a fixed offset, probably in order to achieve similar spatial coverage with the few whiskers they had left. Overall, while freely moving rats used combinations of head and body movements to shift their attentional foci, headfixed rats had to apply a wide range of whisking patterns, varying both amplitudes and offsets, possibly in an attempt to cover as many attentional foci as they could.

Animals use their available resources in an adaptive manner. Focusing on free-air pumps in several contexts, we show here that rats adapt their perceptual behavior to their locomotion plans, both when voluntarily selected and when externally imposed, and suggest that they do so in a way that attempts to optimize coverage of the relevant space and future encounters with objects.

## Materials and Methods

### Whisking in freely-moving rats

The experimental protocol is described in detail in (13). Briefly, the whisking patterns of Wistar strain male albino rats aged 3-6 months were measured (N = 3 for free-air trimmed, N = 4 for free-air single-row). On the day prior to behavioral recording, trimmed whiskers were clipped close to the skin (∼1 mm) under Dormitor anesthesia (0.05 ml/100 g, S.C.). All experimental protocols were approved by the Institutional Animal Care and Use Committee of the Weizmann Institute of Science.

Behavioral experiments were performed in a darkened, quiet room. The behavioral apparatus (See Fig S1) consisted of a holding cage (25 cm width, 35 cm length, 29.5 cm height) with a small door (6.9 cm height, 6 cm width), through which the rats could emerge into the experimental area (18 cm × 20 cm)(49). Both the holding cage and the experimental area were fixed approximately 15 cm above the surface of a table. The experimental area consisted of a back-lit Perspex plate with 1–2 objects (Perspex cubes and cylinders) placed on it. The experimental area was filmed from above by a high-speed, high-resolution camera (1280 × 1024 pix, 500 fps, CL60062, Optronics). An in-house program (E. Segre, Weizmann Institute) triggered the high-speed camera whenever the rat emerged from the holding cage into the experimental area. Video recording stopped when the rat returned to the holding cage.

An experimental session consisted of recording a rat’s whisking behavior whenever the rat was in the experimental area, over a period of 30–120 min. Preceding a session, the animal was placed in the holding cage for a 15-minute acclimation period. During the acclimation period, the door of the holding cage was blocked. The experimental session began with unblocking the door to allow the animal to leave the holding cage and explore the experimental area at will. Each trial started when the rat moved from the holding cage to the experimental area and ended when the rat went back into the holding cage. The length of the experimental session varied depending on the animal’s behavior, and the amount of recorded video. All raw video records were analyzed using the BIOTACT Whisker Tracking Tool http://bwtt.sourceforge.net (50). Protraction angle of all existing macrovibrissa were tracked, as well as the location of both eyes and the tip of the nose. Only tracked segments that did not include object contacts were used for analysis in the current work.

### Whisking in head fixed rats

The experimental protocol is described in details in (23). Briefly, 7 male albino rats were head fixed using screws glued to the skull under anesthesia. After full recovery, rats were gradually adapted to head fixation for 4-5 days. All except for three whiskers (C1, C2, and D1) on either side were clipped close (~1 mm) to the skin during brief (5–10 min) isoflurane anesthesia, and were re-trimmed two to three times a week at least 2 hours before an experiment. A single experiment lasted up to 30 minutes, but terminated earlier if rats showed signs of distress. Each rat had one or two experimental sessions a day, two or three times a week. All experiments were performed in a dark, sound-isolated chamber. Head orientation was estimated by imaging the corneal reflections of two infrared (880 nm) light-emitting diode (LED) spotlights. An imaginary line between the nose and eye on each side of the rat served as a reference line for the whisking angle. Bright-field imaging of the whiskers was accomplished by projecting infrared light (880 nm) with an array of 12×12 LEDs from below the animal. Video acquisition was triggered manually, and high-speed video was buffered and streamed to disks at either 500 or 1,000 frames/s. A total of 255 trials with an average duration of 8.3±3.2 s (mean ± standard deviation, SD) were acquired. 106 “free-air trials” and 149 “contact trials” were intermixed. Only the free-air trials are used for analysis in the current work. Whisker movements were tracked off-line using the *Matlab*-based *WhiskerTracker* image processing software (available at http://code.google.com/p/whiskertracker).

### Analysis

#### Rhythmic decomposition and phase segmentation

All analyses were performed on protraction angle of whisker C2 on the left side of the snout. Analyses of head turns also used whisker C2 on the right side of the snout, and the results from both sides were pooled after mirroring. Signal was first filtered using a low-pass filter (6^th^ order Butterworth, passband cutoff 20 Hz) to remove noise; this filtering was chosen to remove high-frequency noise due to tracking errors, while preserving the pumping information. Whisking phase *φ* was extracted using the Hilbert transform from a high-pass filtered version (cutoff 4 Hz). Peaks and troughs were located within ± *π*/8 of *φ* = 0 and *π*, respectively, and signal was segmented into individual protraction and retraction phases. Offset of each motion was defined as the middle point between trough and peak; amplitude was half the distance between the two points.

#### Pump detection and quantification

Angular trajectory of each phase was differentiated to produce the angular velocity. Retraction velocity profile was negated. A pump was identified whenever the resulting velocity profile has more than one peak. The shapes of individual protraction/retraction phases were previously analyzed in great detail in (20); there, three categories of motion were described: *single pumps*, in which the protraction/retraction velocity profile contains a single peak (these are the ‘default’, unmodulated whisking cycles); *delayed pumps*, in which there are two velocity peaks but there is no reversal in the direction of motion; and *double pumps*, where motion in the opposite direction (e.g., backward during protraction) is detected.

We quantify the *pump strength σ*_*pump*_ by using the formula: 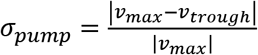, where *v*_*max*_ is the maximal velocity during the protraction/retraction and *v*_*through*_ the velocity at the lowest trough. Note that by this definition *σ*_*pump*_ = 0 for ‘single pump’ (i.e., no pump), 0 < *σ*_*pump*_ < 1 for ‘delayed pump’ and *σ*_*pump*_ > 1 for double pump’. We did not observe any discontinuity in the distributions around *σ*_*pump*_=1, and therefore delayed and double pump profiles were lumped together; we refer to these two profiles simply as *pumps*.

#### Head motion

Total head translational velocity in freely moving experiments was defined as the velocity of the middle point in between the two eyes (see Fig S3, panel A). Head direction was defined as the direction of the line connecting this point and the tip of the nose. Length of this line was defined as head size used to normalize translational velocity (to units of heads/s). Direction of translational velocity was than subtracted from the head direction to obtain the translational direction in head-centered coordinates. The projection of this vector on the line pointing towards the nose was defined as thrust (longitudinal translational velocity), while the orthogonal projection was defined as slip (transverse translational velocity).

#### Statistics

Random permutations were used to evaluate significance of correlations (Figs 2A, 3 and 4) and run length distributions (Fig 2B); the bootstrap method was used in all comparisons between sets (Figs 5, 6D and 7). Unless stated otherwise, 5000 permutations/draws were used in each comparison. When comparing offset and amplitude distributions in different contexts (Figs 1B, 7), the samples of each set are not statistically independent due to temporal correlations (21, 23). Therefore, for each segment analyzed we computed the autocorrelation functions of these variables and measured the lag (number of cycles) at which their significance dropped below the p=0.05 level (using random permutations as control). The amplitude/offset sequence was then diluted by this lag to obtain statistically independent samples.

#### Whisking envelope entropy

To estimate the dispersion of the whisking envelope (Fig 6D), we measured the bivariate probability distribution *p*(*θ*_*amp*_,*θ*_*off*_), by binning each variable into 25 bins. We then measured the information entropy using the formula: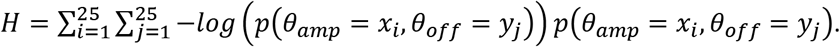. To enable statistical comparison between contexts random sets of identical size from all datasets were required. 75 different collections of tracked segments were randomly generated from each dataset so that the total duration of each collection was approximately 30% of the total length of the smallest dataset. Entropy was then calculated for each of the resulting subsets to produce a distribution of entropies for each dataset (boxplots Fig 6D). Entropy was also calculated for each of the datasets in its entirety (inset in Fig 6D).

## Acknowledgements

We thank Sahar Froim for experimental assistance and Mitra Hartmann and Mathew Diamond for discussions and advices. This project has received funding from the Israel Science Foundation (grant No. 1127/14), the Minerva Foundation funded by the Federal German Ministry for Education and Research, the Israel Ministry of Defense, and the United States-Israel Binational Science Foundation (BSF, grant No. 2017216). E.A. holds the Helen Diller Family Professorial Chair of Neurobiology.

---

**Fig S1:**
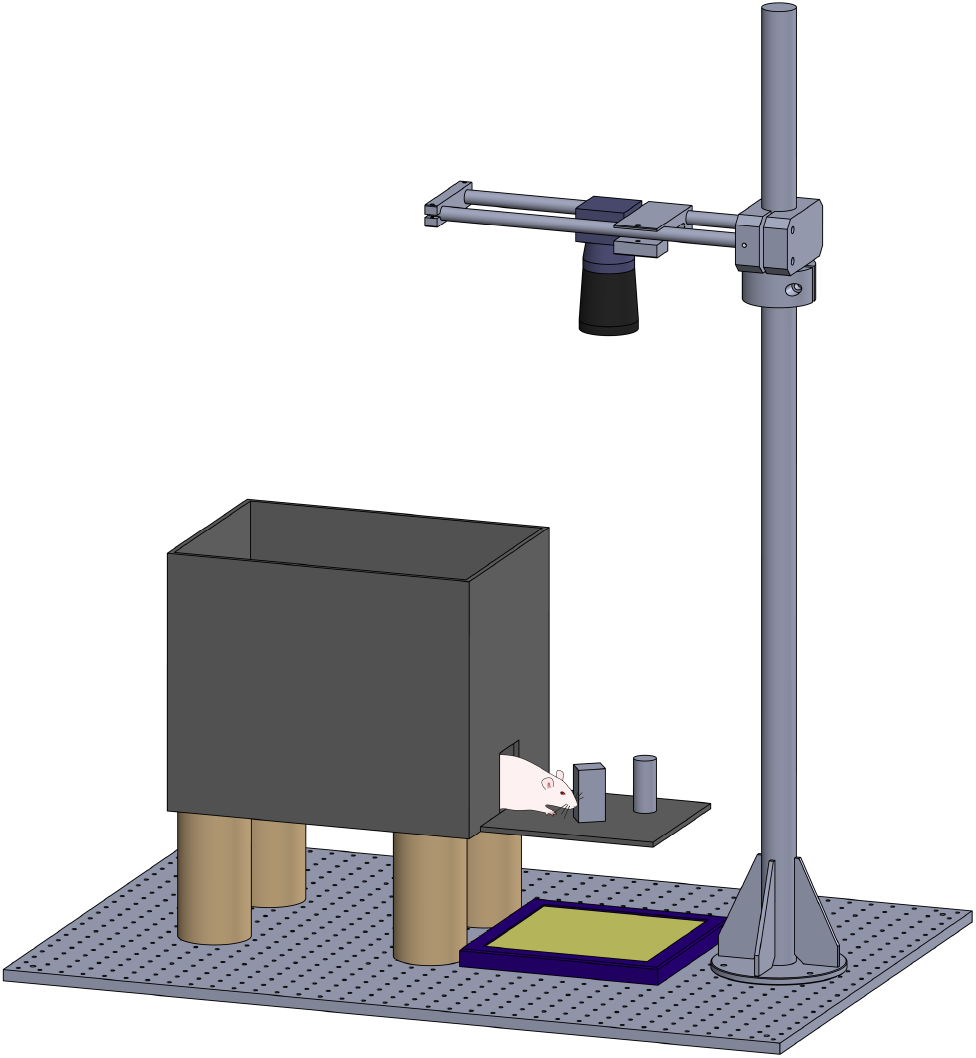
Freely moving apparatus. The apparatus included: a holding cage with door (left) approximately 15 cm above the surface of a table; the experimental area (back-lit Perspex plate with acrylic cubes and cylinders); a high-speed, high-resolution overhead camera.

**Fig S2:**
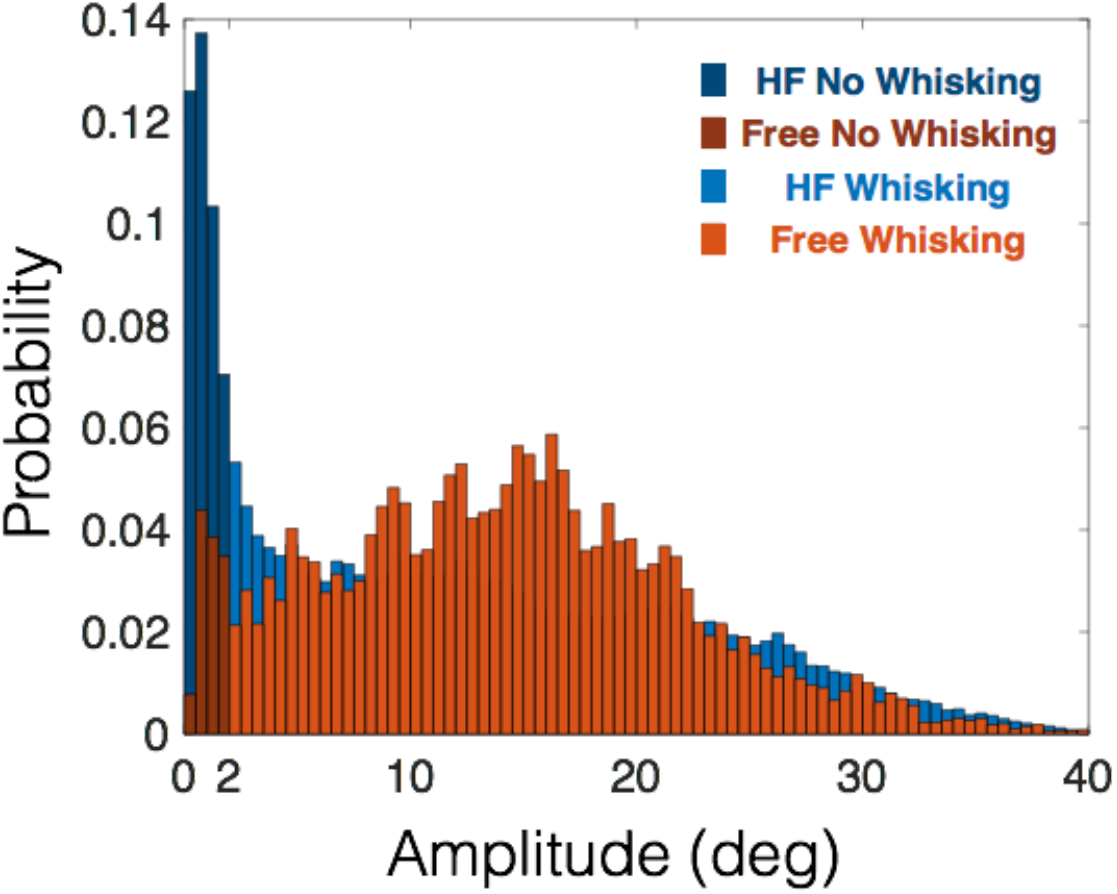
Whisking amplitude distributions and whisking threshold. Probability distributions of whisking amplitude, as computed by rhythmic decomposition, for headfixed (blue) and freely moving (red) trimmed rats. The noticeable peak at extremely low amplitudes (<2 deg, dark colors) relates to periods of no whisking.

**Fig S3:**
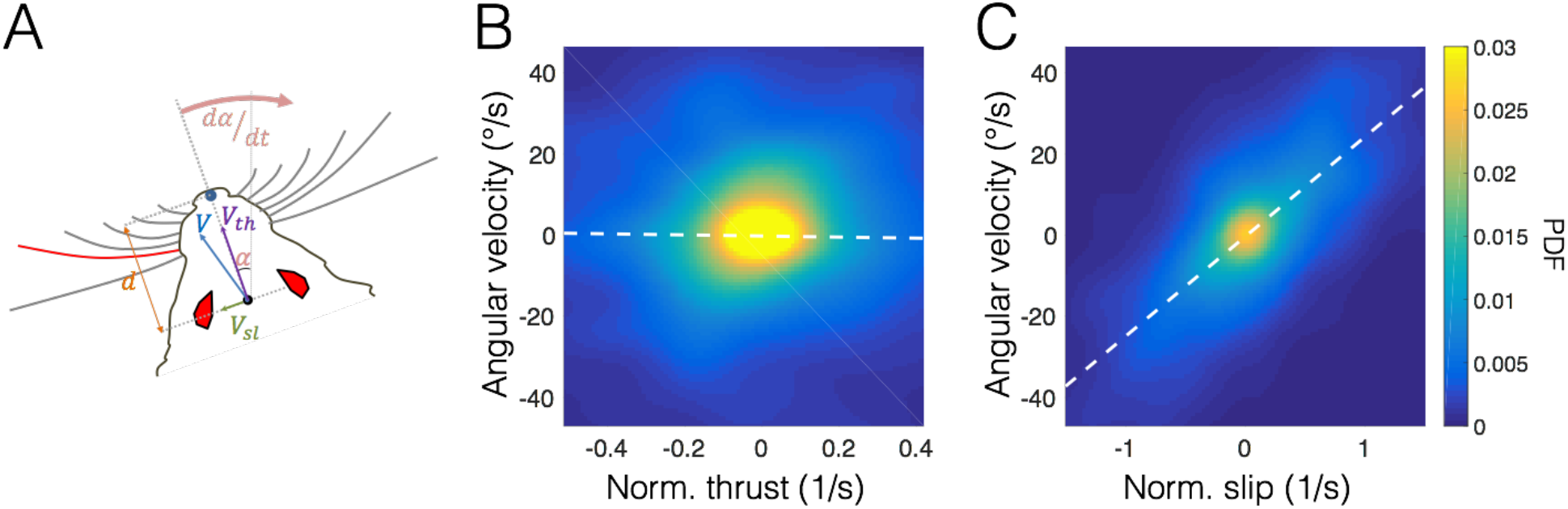
Head-motion analysis. **(A)** Head tracking scheme. The two eyes and the tip of the nose were tracked in each video frame. Midpoint between eyes was defined as the head center. The azimuth of the line connecting this point and the nose is the head direction *α*, while the length of this line is the head size *d*. The time derivative of the head direction, *dα*/*dt*, is defined as turn (head rotation). The time derivative of the head location is the head velocity *V*, which has a longitudinal component thrust (*V*_*th*_) and a transverse component termed slip (*V*_*sl*_). These components were normalized to the head size *d* and so are presented in units of heads/s. **(B)** Joint probability density of the thrust and turn variables. Pearson coefficient R=−0.035. Dashed white line: linear regression. **(C)** Joint probability density of the slip and turn variables. Pearson coefficient R=0.863. Dashed white line: linear regression.

**Fig S4:**
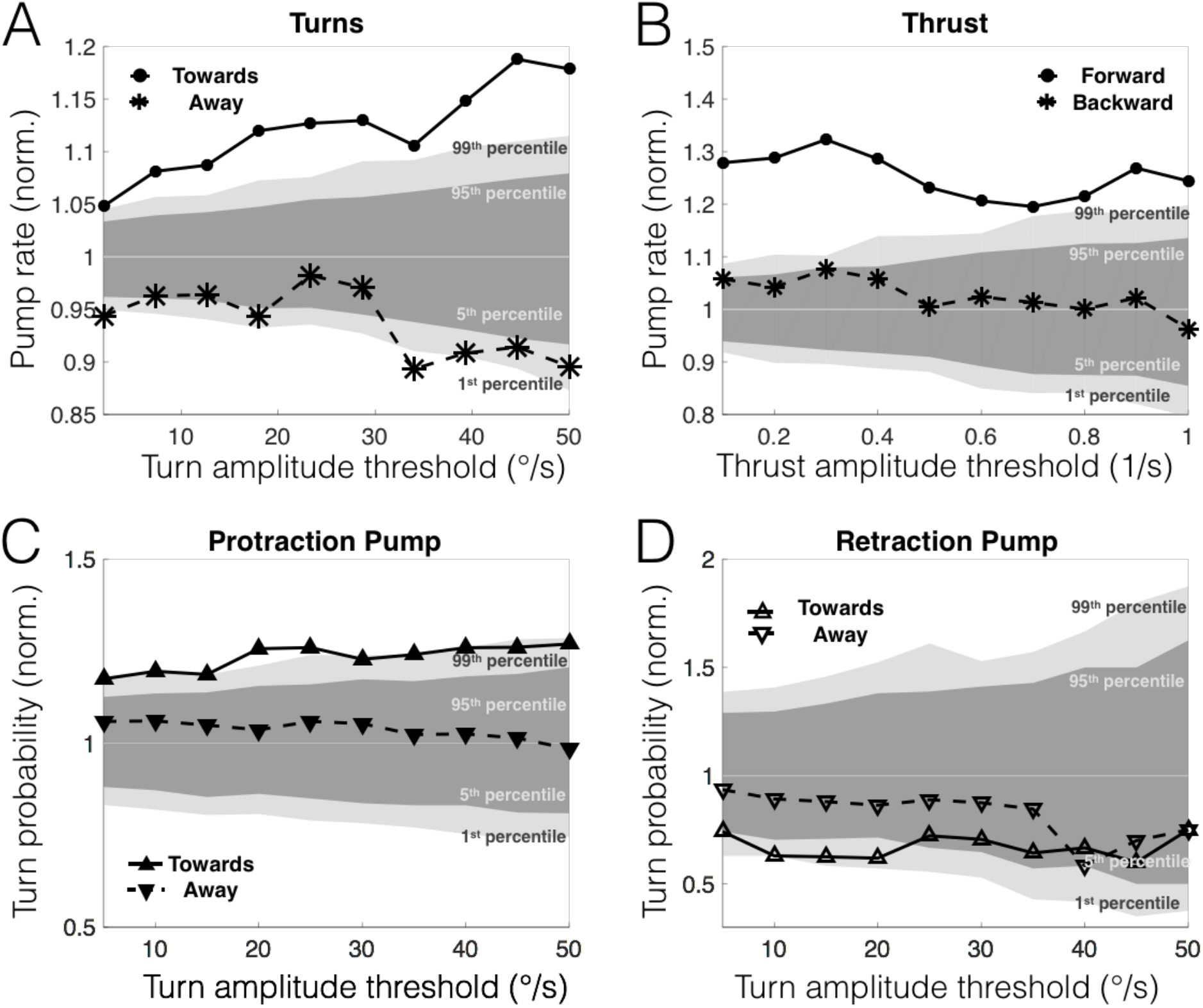
Robustness of pump-rate indication prior to change of motion. Abscissa in all panels: threshold of used to identify individual motions (turn/thrust). Grey shadings in all panels: distributions of random-permutation generated controls. **(A-B)** The protraction pump rate (normalized to median control levels) 100 ms prior to change of motion, computed for various levels of detection threshold. **(A)** Turn. Solid line with filled circles: towards pumping side; dashed line with asterisks: away from pumping side. Elevation in pump probability prior to turning towards pump is significant throughout the threshold range. **(B)** Thrust. Solid line with filled circles: onset of motion forward; dashed line with asterisks: onset of motion backwards. Elevation in probability of forward onset is significant throughout the threshold range. **(C-D)** The turn probability (normalized to median control levels) 100 ms following a pump, computed for various levels of detection threshold. Solid line with triangles: towards pumping side; dashed line with upside-down triangles: away from pumping side. **(C)** Protraction pump. Elevation in probability of turning towards pump is significant throughout the threshold range. **(D)** Retraction pump.

